# Simulations Predict Stronger CRISPRi Transcriptional Repression in Plants for Identical than Heterogeneous gRNA Target Sites

**DOI:** 10.1101/2024.04.22.590637

**Authors:** Helen Scott, Alessandro Occhialini, Scott C. Lenaghan, Jacob Beal

## Abstract

Plant synthetic biologists have been working to adapt the CRISPRa and CRISPRi promoter regulation methods for applications such as improving crops or installing other valuable pathways. With other organisms, strong transcriptional control has typically required multiple gRNA target sites, which poses a critical engineering choice between heterogeneous sites, which allow each gRNA to target existing locations in a promoter, and identical sites, which typically require modification of the promoter. Here, we investigate the consequences of this choice for CRISPRi plant promoter regulation via simulation-based analysis, using model parameters based on single gRNA regulation and constitutive promoters in *N. benthamiana*. Using models of 2 to 6 gRNA target sites to compare heterogeneous versus identical sites for tunability, sensitivity to parameter values, and sensitivity to cell-to-cell variation, we find that identical gRNA target sites are predicted to yield far more effective transcriptional repression than heterogeneous sites.

## 1 Introduction

Genetically engineered plants hold immense promise for helping to address critical global challenges such as food security, environmental sustainability, and renewable resource production. Establishing engineering control over a transcriptional regulation has been an important tool for other organisms and is expected to be so in plants as well, which is inspiring many efforts in this area (e.g., [1, 2, 3, 4, 5]).

Two of the key properties for effective promoter regulation are orthogonality and high dynamic range. Orthogonality refers to the promoter’s ability to function independently of the plant’s native regulatory elements. This isolation is beneficial to both the host plant and the engineered circuit [1]. By using orthogonal promoters, the circuit’s desired function is protected from perturbations caused by the host plant, and the host plant itself is protected from undesired regulatory impacts from the circuit [6]. Dynamic range refers to the range of output expression levels (e.g., RNA or protein production) that is achieved when the promoter is in its binary “on” state (high rate of transcription) or “off” state (low rate of transcription). A promoter with a high dynamic range is capable of producing a significant difference in expression levels between the on and off states, both allowing greater modification of behavior and enabling more complex regulatory networks [7].

CRISPR/Cas transcriptional regulatory systems are both orthogonal in plants and have the potential for a high dynamic range. In these systems, the catalytically inactivated Cas9 protein (dCas9), either alone or fused to a transcriptional activation or repression domain, binds to a specific promoter region determined by the sequence of a customized guide RNA (gRNA) [8], thereby activating (CRISPRa) or inhibiting (CRISPRi) transcription. Both have previously been used in plants, but have often resulted in a low dynamic range in both plants [9, 10, 11] and other systems [12, 13, 14]. However, the efficacy of regulation can be increased by using multiple gRNA target sites to control a single promoter. For example, previous studies have used 3-4 gRNAs for CRISPRa in mammalian cells [15], 2 gRNAs for CRISPRi in mammalian cells [16], and 4 gRNAs for CRISPRi in yeast [17]. Multiple gRNAs have also been shown to have synergistic effect for CRISPRa in plants [11], but have not yet achieved the dynamic range seen in other organisms.

A key question when using gRNA to target multiple sites on a promoter, is whether to have the targets be heterogeneous (i.e., each gRNA has a unique sequence) or identical (i.e., the same gRNA is used for each target site). Heterogeneous gRNA target sites are often easier to use, since there is no need to find or engineer identical nearby sequences, but each site requires a separate dCas9 binding events to contribute to transcriptional repression (Figure 1A). On the other hand, using multiple identical binding sites reduces the competition for Cas9 between gRNA species, and can allow for a single binding event to result in multiple target sites being occupied due to the dCas9-gRNA complex laterally diffusing along the DNA strand (Figure 1B) rather than unbinding completely [18]. Heterogeneous target sites have been used in mammalian cells [15], and yeast [17], and identical target sites in mammalian cells [16]. However, in these cases, the choice between heterogenous and identical target sites was a pragmatic choice based on the ease of engineering. To the best of our knowledge, there has not been a detailed analysis of the efficacy of heterogenous or identical target sites for CRISPRa or CRISPRi in any system.

**Figure 1:**
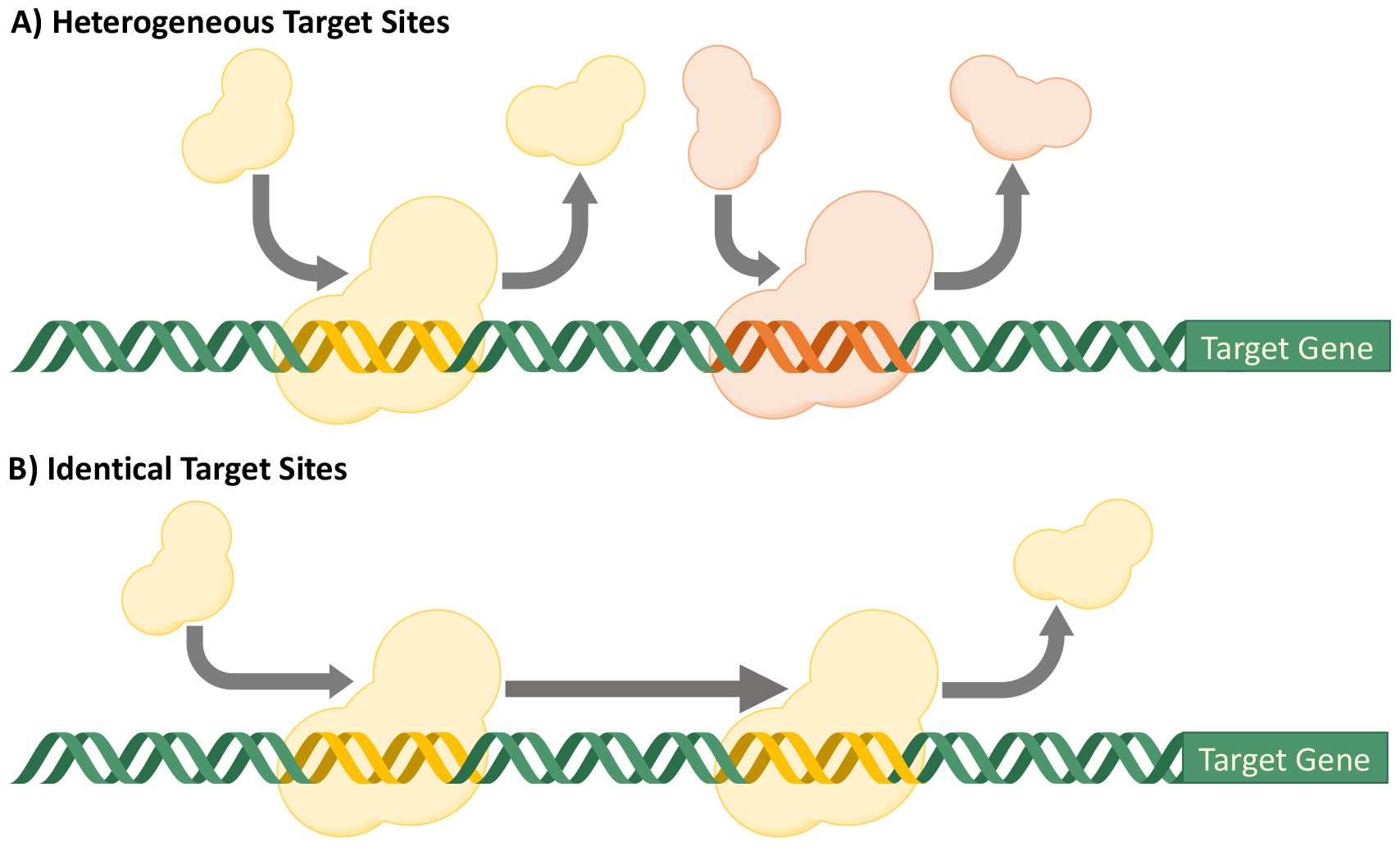
Heterogeneous vs. identical gRNA gRNA target site strategies. A) Heterogeneous gRNA target sites are easier to target, since there is no need to find or engineer identical nearby sequences, but each site requires a separate dCas9 binding event to contribute to transcriptional repression. B) Identical gRNA target sites avoid competition for Cas9 between gRNA species and also allow for a greater probability of the second binding site being occupied, as the dCas9-gRNA complex can “scan” to occupy the second site after unbounding from the first site.

To better understand the tradeoffs for CRISPRi between heterogeneous and identical gRNA target sites for CRISPRi in plants, we thus perform a simulation-based analysis of the potential for building highly repressible synthetic plant promoters using multiple gRNA targets. Specifically, we built a model of CRISPRi regulation in *N. benthamiana* using parameters drawn from single gRNA regulation and constitutive promoters in that organism and evaluate it for up to six gRNA heterogeneous or identical targets. Evaluating these circuits in simulation with respect to a range of biologically plausible parameters, we find a strong benefit for using identical gRNA target sites, both in achievable fold repression and in the scaling of repression per additional gRNA target site.

## 2 Results

We first detail the different repressible promoter architectures under investigation, then present an exploration of their behaviors with respect to biologically plausible parameters based on studies of single gRNA regulation and constitutive promoters in *N. benthamiana*. Further evaluation of the promoters’ tunability, sensitivity to parameter values, and sensitivity to cell-to-cell variation (presented in the Supporting Information) shows that the conclusions from this study are not sensitive to the specific values used in our investigation.

### 2.1 Generation of Possible Promoter Architectures

A simple CRISPRi system with a single gRNA and single target site is shown in Figure 2A. The system consists of an engineered genetic vector (V1) for expression of the single guide RNA (sgRNA1) and a second vector (V2) for expression of dCas9 and the GFP fluorescent marker controlled by the target promoter sequence.

**Figure 2:**
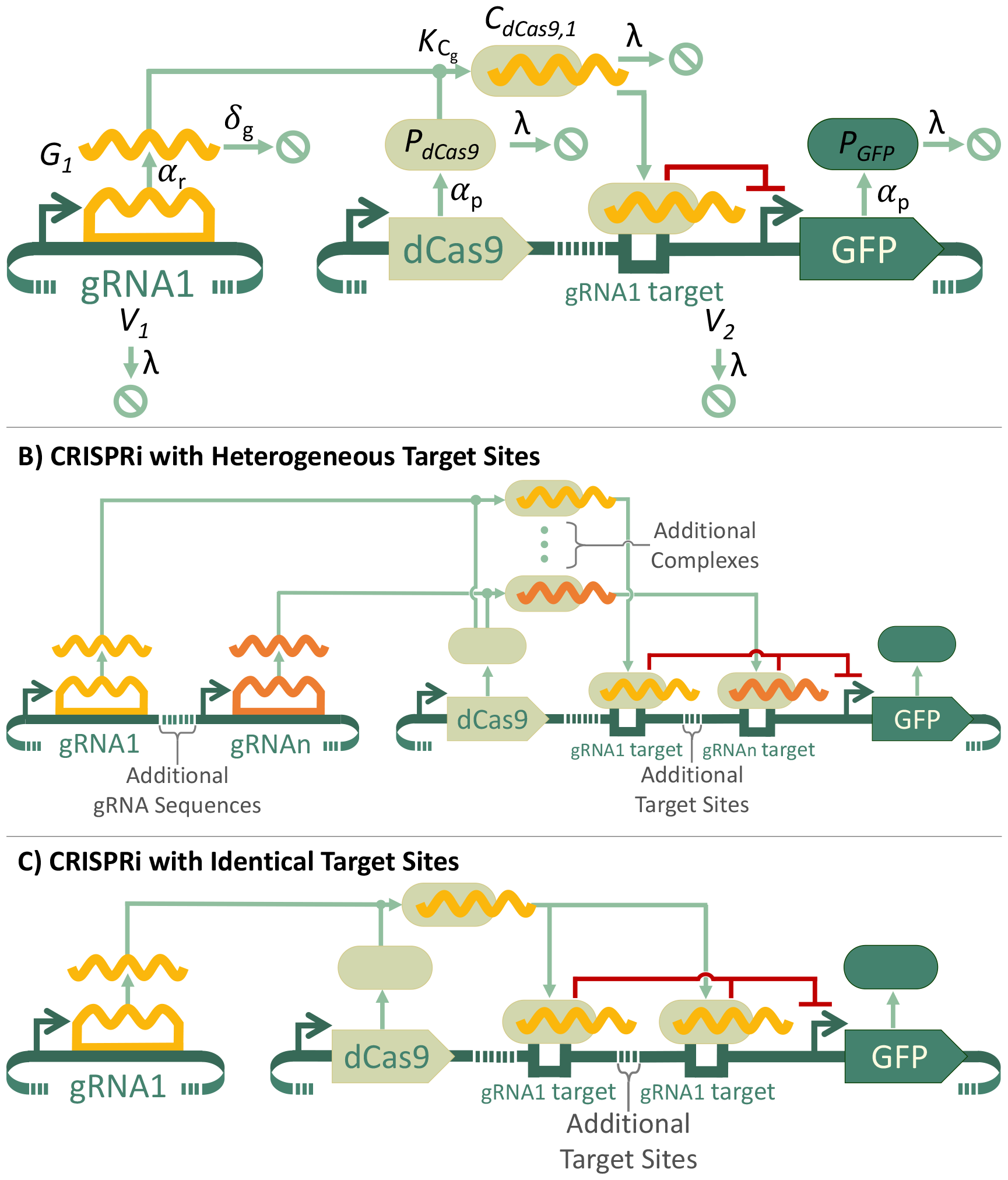
Circuit diagrams for different CRISPRi architectures. A) Base CRISPRi architecture, where a single gRNA sequence binds to a single target site in the GFP promoter, including mathematical symbols for species and process rates. In the “off” state, the gRNA vector is included and represses GFP expression, while in the “on” state the gRNA vector is omitted (*V*_1_ = 0) and GFP is not repressed. B) Generalized system for CRISPRi with heterogeneous target sites, numbered 1 through *n*. C) Generalized system for CRISPRi with identical target sites, numbered 1 through *n*. All circuits are shown using SBOL visual notation [19].

The utilization of a two-component strategy has several advantages. The control of multiple transgenes from individual expression cassettes integrated in the same transformation vector is often difficult. The level of transgene expression can be affected by both direct acting regulatory elements and proximal regulatory elements located in other transgene cassettes, causing different expression levels from the same transgene cassette based on vector context. Furthermore, the utilization of a two-component strategy allows improved system flexibility by enabling “mix-and-match” combination of different components. The utilization of separate vectors for sgRNA and dCas9 expression also enables modulation of the stoichiometry of the two components simply by changing vector dosages during transformation of plant cells.

The expression of dCas9 is constitutive, and the dCas9 protein can bind to the gRNA to form a dCas9-gRNA complex (in this case dCas9-gRNA1), which in turn binds to a gRNA binding site in the GFP promoter that represses transcription. Figure 2A also shows the mathematical symbols for the concentrations of the molecular species in the model and the parameterized rates that modulate the changes in these concentrations. All of the stable molecules, including the genetic constructs and the proteins, dilute at a rate of *λ*, while the gRNA molecules degrades at a faster rate of *δ*_*g*_. Each genetic product is associated with its own production rate, *α*, where *α*_*r*_ is a transcription rate to produce gRNA and *α*_*p*_ is a combined transcription and translation rate to produce protein. The different *α* rates can be varied independently to reflect the strength of each protein’s individual promoter. The dynamics of the dCas9-gRNA complex formation are described by the binding constant *K*_*C*_*g*. An ordinary differential equation model was constructed for this system and for constitutive expression in the absence of gRNA (the “on” condition) along with a set of biological plausible base parameter values. Details for equation and parameter development are described in the Methods and the equations and parameter values are provided in the Supporting information.

To investigate the potential for stronger repression, we generated models for using 2, 3, 4, 5, or 6 heterogeneous or identical gRNA target sites in a promoter. The generalized forms of the heterogeneous and identical target site systems are shown in Figure 2B-C. As the focus of this investigation is the regulatory effects of multiple gRNA target sites, we defer any additional details related to construct ordering and specific gRNA site location for future investigation. For purposes of this discussion, then, the ordering of functional units and the placement of gRNA target sites are both notional, selected primarily for clarity of illustration. Ordinary differential equation models were generated for these 10 multiple gRNA target circuits using the same methods as for the single gRNA target circuit. These models are also provided in the Supporting Information and use the same parameter values as the single target circuit.

### 2.2 Analysis of Promoter Repressibility

We use simulation to evaluate the strength of repression seen for each of the 11 promoter architectures under consideration. Figure 3 shows the results of simulating all possible promoter architectures over a 200 hour time course using the base parameter values, starting with an initial 10 copies of the genetic construct encoding the gRNA(s), and 3 copies of the genetic construct with dCas9 and GFP. Figure 3A shows the concentration of the GFP over the time course for each of the possible promoter architectures. The fold repression can be calculated at any point for one of the models using the ratio between the Base Expression (no gRNA) model and the model of interest. Figure 3B shows the average fold repression for each of the models between 72 and 96 hours of the time course. The mean fold repression was shown for the 72-96 hour window because during this day GFP expression is no longer rising in the unrepressed system (Base Expression, No Regulation) and vector dilution has not yet significantly decreased levels of dCas9-mediated repression.

**Figure 3:**
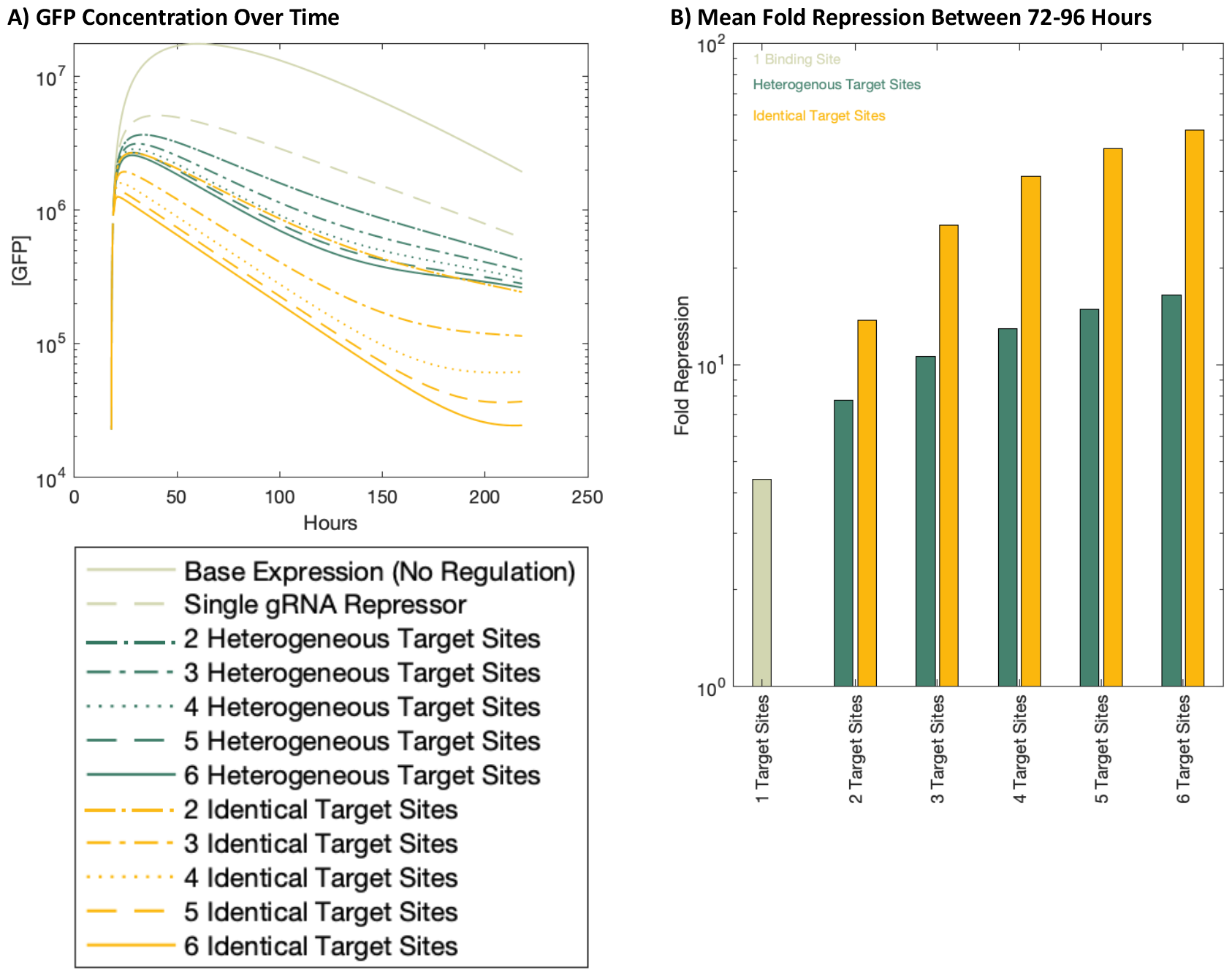
Simulation of GFP production for all promoter architectures using base parameter values. A) The concentration of GFP over time for each system. The promoter architectures are grouped by color for the different gRNA target site strategies, with dark green lines representing all heterogeneous target site systems, and gold lines representing identical target site systems. Pale green lines represent the systems with no gRNA control and with a single gRNA target site. B) The fold repression for each promoter architecture, comparing the GFP concentrations shown in panel A for each repressed system against the Base Expression (No Regulation) system. Bar height indicates the mean fold repression between 72 and 96 hours in the time course.

Figure 3 shows higher levels of repression when using identical target sites than using the same number of heterogeneous target sites. This is seen in Figure 3A, where the GFP concentrations for the identical target site repression systems (yellow lines) are lower than almost all GFP concentrations for the heterogeneous target site repression systems (dark green lines), and in Figure 3B, where the fold repression for identical target sites (yellow bar) is always higher than the fold repression for the same number of heterogeneous target sites (dark green bars). Figure 3B shows that this increased efficacy in repression allows fewer identical target sites to have the same effect as a larger number of heterogeneous target sites. For example, two identical gRNA target sites gives roughly equivalent levels of repression as five heterogeneous gRNA target sites.

Figure 3 also shows that increasing the number of gRNA target sites in a promoter increases the ability to repress that promoter. However, adding identical target sites increases that ability more quickly. For example, when adding a sixth identical target site mean repression goes from approximately 47-fold with five identical target sites, to 53-fold repression with the sixth. This difference of over 6-fold repression is much larger than the equivalent 1.5-fold difference between five and six heterogeneous target sites. For both the heterogeneous and identical target sites, Figure 3 shows that the largest increase in repressibility comes from adding a second target site to the single gRNA system. Adding a third, fourth, fifth, or sixth binding site, increases repression, but with a smaller increase each time.

Finally, while models with low levels of repression (such as the Single gRNA model) result in an approximately log-linear decrease in GFP concentration, other models show inflections in the rate of change in GFP. This inflection is both stronger and begins later for models with higher levels of repression, with the maximum found in the six identical target site model (Figure 3A).

## 3 Discussion

Models with identical gRNA target sites show greater repression than models with the same number of heterogeneous target sites, and additional identical target sites caused a greater increase in repression than the addition of heterogeneous target sites. This difference between the heterogeneous target sites and identical target sites is caused by introducing dependence between the effect of the multiple gRNA target sites. Heterogeneous target sites have independent effects; for each target site to have an effect on repression, the dCas9 must bind to the correct site on DNA, which has a relatively low likelihood of occurring. While, the initial binding of a dCas9-gRNA complex to the DNA is still relatively low for a promoter with multiple closely spaced identical gRNA sites, there is now a higher likelihood of the dCas9 complex remaining attached, as Cas9 does not remain strongly bound to a single site, but instead scans short distances along DNA for additional target sequences [18]. By shifting from events with all low independent likelihoods of occurring to one low likelihood event of binding, and subsequent higher likelihood events of binding, multiple identical target sites are expected to yield higher levels of transcriptional repression. The Supporting Information includes parameter perturbation studies that show that these conclusions are not dependent on precise parameter values.

Figure 3A also shows two phenomena that we would expect based on the biophysical constraints on the system. The first is a decrease in GFP for the “Base Expression (No Regulation)” system. While there is no repressor turning off GFP expression, protein levels still go down in transient expression, as the GFP expression construct (*V*_2_) dilutes in actively dividing cells. A second phenomenon seen is a plateaued level of GFP for the very strong repressors (e.g. “6 Identical Target Sites”), as a tight off signal will eventually not be able to go significantly lower.

While this model does not account for the physical design of the promoter, including the precise location of the gRNA target sites, recreating these results experimentally will depend on the physical constrains of dCas9 lateral diffusion. Lateral diffusion of Cas9 is typically limited to local distance of approximately 20 basepairs [18] and is facilitated by binding to protospacer adjacent motif (PAM) sequences. To account for these limitations, promoter design could insert PAM sites within approximately 20 basepairs of each other, potentially including PAM sites that are not adjacent to a target site, in order to prolong lateral diffusion between targets. These design considerations will need to be balanced with the needs of stability and functionality of the promoter.

For further development of identical target site promoters, next steps include more accurate simulation of repression with models of the biophysical dynamics of gRNA and promoter DNA binding, as well as validation of these circuits in the laboratory. If the expected high levels of repression are verified experimentally, CRISPRi with identical target sites may be a valuable addition to the plant synthetic biology tool kit, with potential applications in food security, environmental monitoring, and biomaterial production.

## 4 Methods

### 4.1 Model Construction

All of the models investigated were represented using SBOL3 [20, 21]. Specifically, we constructed SBOL generators for each of the modular elements of the circuit: the expression of dCas9 and gRNA(s), their binding kinetics, and the transcriptional regulation of GFP. Because of the modular creation of the circuits, the number of gRNAs and the number of interactions between each gRNA and the GFP promoter could be varied combinatorially to generate a genetic regulatory network model for each of the possible configurations. Both LaTeX equations and MATLAB code were then programmatically generated for each genetic regulatory network. Complete information about the models used, including full ODE equations for all systems, is provided in the Supporting Information. The SBOL files for all circuit designs, as well as MATLAB simulation files are provided in the Supporting Information. Model generators and products are also available on GitHub at https://github.com/TASBE/CRISPRi-promoters.

### 4.2 Parameter Fitting

As detailed in the Supporting Information, the parameters needed for the models were taken from previously published studies, fit to a long time course experiment, or hypothesized based on observed repression values. For parameters fit to newly generated experimental data, we performed a least-squares fit for parameters in a logarithmic parameter value space in MATLAB using the ODE model for no gRNA transcriptional repression. The complete set of parameter base values and their sources is shown in the Supporting Information.

## 5 Acknowledgments

This work was supported by DARPA contract HR0011-18-2-0049. This document does not contain technology or technical data controlled under either U.S. International Traffic in Arms Regulation or U.S. Export Administration Regulations. Views, opinions, and/or findings expressed are those of the author(s) and should not be interpreted as representing the official views or policies of the Department of Defense or the U.S. Government. Approved for public release, distribution unlimited (DISTAR Case 39302).

## References

[1] Jenny Koukara and Kalliope K Papadopoulou. “Advances in plant synthetic biology approaches to control expression of gene circuits”. en. In: Biochem. Biophys. Res. Commun. 654 (Apr. 2023), pp. 55–61.

[2] Marta Vazquez-Vilar, Sara Selma, and Diego Orzaez. “The design of synthetic gene circuits in plants: new components, old challenges”. en. In: J. Exp. Bot. 74.13 (July 2023), pp. 3791–3805.

[3] Alexander C Pfotenhauer et al. “Building the Plant SynBio Toolbox through Combinatorial Analysis of DNA Regulatory Elements”. en. In: ACS Synth. Biol. 11.8 (Aug. 2022), pp. 2741–2755.

[4] Jennifer A N Brophy et al. “Synthetic genetic circuits as a means of reprogramming plant roots”. en. In: Science 377.6607 (Aug. 2022), pp. 747–751. ISSN: 0036-8075, 1095-9203. doi: 10.1126/science.abo4326. URL: http://dx.doi.org/10.1126/science.abo4326.

[5] James P B Lloyd et al. “Synthetic memory circuits for stable cell reprogramming in plants”. en. In: Nat. Biotechnol. 40.12 (Dec. 2022), pp. 1862–1872. ISSN: 1087-0156, 1546-1696. doi: 10.1038/s41587-022-01383-2. URL: http://dx.doi.org/10.1038/s41587-022-01383-2.

[6] Stephanie C Amack and Mauricio S Antunes. “CaMV35S promoter – A plant biology and biotechnology workhorse in the era of synthetic biology”. In: Current Plant Biology 24 (Dec. 2020), p. 100179.

[7] Jacob Beal et al. “Meeting Measurement Precision Requirements for Effective Engineering of Genetic Regulatory Networks”. In: ACS Synthetic Biology 11 (3 Mar. 2022), pp. 1196–1207.

[8] Lei S Qi et al. “Repurposing CRISPR as an RNA-guided platform for sequence-specific control of gene expression”. en. In: Cell 152.5 (Feb. 2013), pp. 1173–1183.

[9] Levi G Lowder et al. “A CRISPR/Cas9 Toolbox for Multiplexed Plant Genome Editing and Transcriptional Regulation”. en. In: Plant Physiol. 169.2 (Oct. 2015), pp. 971–985.

[10] Camilo Calvache et al. “Strong and tunable anti-CRISPR/Cas activities in plants”. en. In: Plant Biotechnol. J. 20.2 (Feb. 2022), pp. 399–408.

[11] Agnieszka Piatek et al. “RNA-guided transcriptional regulation in planta via synthetic dCas9-based transcription factors”. en. In: Plant Biotechnol. J. 13.4 (May 2015), pp. 578–589.

[12] Nader Alerasool et al. “An efficient KRAB domain for CRISPRi applications in human cells”. en. In: Nat. Methods 17.11 (Nov. 2020), pp. 1093–1096.

[13] Brady F Cress et al. “CRISPRi-mediated metabolic engineering of E. coli for O-methylated anthocyanin production”. en. In: Microb. Cell Fact. 16.1 (Jan. 2017), p. 10.

[14] Rui Miao et al. “CRISPR interference screens reveal growth-robustness tradeoffs in Synechocystis sp. PCC 6803 across growth conditions”. en. In: Plant Cell (July 2023).

[15] Albert W Cheng et al. “Multiplexed activation of endogenous genes by CRISPR-on, an RNA-guided transcriptional activator system”. en. In: Cell Res. 23.10 (Oct. 2013), pp. 1163–1171.

[16] Samira Kiani et al. “CRISPR transcriptional repression devices and layered circuits in mammalian cells”. In: Nature Methods 11 (7 July 2014), pp. 723–726.

[17] Nicholas S McCarty et al. “Rapid assembly of gRNA arrays via modular cloning in yeast”. en. In: ACS Synth. Biol. 8.4 (Apr. 2019), pp. 906–910.

[18] Viktorija Globyte et al. “CRISPR /Cas9 searches for a protospacer adjacent motif by lateral diffusion”. en. In: EMBO J. 38.4 (Feb. 2019). ISSN: 0261-4189, 1460-2075. doi: 10.15252/embj.201899466. URL: http://dx.doi.org/10.15252/embj.201899466.

[19] Hasan Baig et al. “Synthetic biology open language visual (SBOL visual) version 3.0”. en. In: J. Integr. Bioinform. 18.3 (Oct. 2021).

[20] James Alastair McLaughlin et al. “The Synthetic Biology Open Language (SBOL) Version 3: Simplified Data Exchange for Bioengineering”. en. In: Front Bioeng Biotechnol 8 (Sept. 2020), p. 1009.

[21] Lukas Buecherl et al. “Synthetic biology open language (SBOL) version 3.1.0”. In: Journal of Integrative Bioinformatics 20.1 (2023), p. 20220058. doi: 10.1515/jib-2022-0058. URL: https://doi.org/10.1515/jib-2022-0058.

