## Supplementary S1 for "Simulations Predict Stronger CRISPRi Transcriptional Repression in Plants for Identical than Heterogeneous gRNA Target Sites"

### Supplementary S1: Derivation of Equations and Parameter Perturbation for CRISPRi Gene Expression Repression

Helen Scott[1], Alessandro Occhialini[2,3], Scott C. Lenaghan[3,4], and Jacob Beal[1]

[1]Intelligent Software and Systems, RTX BBN Technologies, Cambridge, MA 02138, USA

[2]Department of Plant Sciences, University of Tennessee, 112 Plant Biotechnology Building 2505 E J Chapman Drive, Knoxville, TN 37996, USA

[3]Center for Agricultural Synthetic Biology (CASB), University of Tennessee, 2640 Morgan Circle Dr., Knoxville, TN 37996, USA

[4]Department of Food Science, University of Tennessee Institute of Agriculture (UTIA), 102 Food Safety and Processing Building 2600 River Dr., Knoxville TN 37996, USA

April 22, 2024

### 1 Organization

This document presents the equations used for modeling the various CRISPRi promoter architectures. Section 2 discusses the derivation of the differential equation models used for each individual component in the circuits. Section 3 then composes these elements into all of the promoter architectures investigated.

Section 4 identifies the parameter values used in the models and the sources for those parameters. Section 5 then analyzes the sensitivity of the model to those parameter values.

Table 1 presents the variables used throughout this document, using the following typical conventions:

- Molecule counts are upper case, with the class of molecule indicated by latter (e.g., $G$ for gRNA, $P$ for protein)
- Particular species are indicated by subscript (e.g., $G_i$ indicates a specific gRNA).
- Rate constants are lower case Greek, except $k$ for reaction rate constants.
- Hill-Langmuir equations use $n$ for coefficient and $K_R$ for dissociation constants.

### 2 Model Construction

The possible repression circuits considered here are made of several core biological mechanisms and repeating control mechanisms. We model each mechanism with a standard differential equation model, selected for modularity and abstraction in order to keep the number of required parameters as low as possible. The mechanisms considered are: genetic production by transcription and translation from the engineered vector and the degradation or dilution of products over time, CRISPR complex formation and, and transcription factor regulation of expression.

#### 2.1 Core Biological Mechanisms: Transcription, Translation, Degradation, and Dilution

We consider two classes of products, gRNA and proteins. For a gRNA species $G_i$, its production depends only on the rate of transcription from copies of the vecotr. Decay is assumed to be the standard exponential and dilution negligible (i.e., non-dividing cells). This can be expressed as:

$$\frac{\mathrm{d}[G_i]}{\mathrm{d}t} = \alpha_{r,i} V - \delta_g [G_i] \tag{1}$$

Where $G_i$ is the the concentration of the RNA product, $\alpha_{r,i}$ the transcription rate of the controlling promoter in the absence of regulation, and $\delta_g$ the degradation rate for unbound gRNA. All symbols are also defined in Table 1.

| Variable | Meaning |
|---|---|
| $V_i$ | Construct vector, species i |
| $G_i$ | gRNA, species i |
| $P_{\text{dCas9}}$ | dCas9 protein species |
| $P_{\text{GFP}}$ | GFP protein species |
| $C_{\text{dCas9},i}$ | Complex of dCas9 protein and gRNAi |
| $\alpha_{r,i}$ | Transcription rate, species i |
| $\alpha_{p,i}$ | Coupled transcription and translation rate, species i |
| $\delta_g$ | gRNA degradation rate |
| $\lambda$ | Stable Molecule Dilution Rate |
| $k_{C_g}$ | dCas9 and gRNA binding rate |
| $n$ | Hill coefficient |
| $K_A$ | Hill Activation Constant |
| $K_R$ | Hill Repression Constant |
| $m$ | Number of Identical Binding Sites |

Table 1: Table of variable symbols and meanings

For protein products, the rate of formation of the product is dependent on both transcription and translation. Transcription and translation are modeled as a joint rate in the model under the assumption that mRNA decay rate is fast compared to decay rate of the proteins under consideration, which allows the intermediate stage of mRNA to be abstracted away, reducing the number of parameters. Production and decay of a protein can thus be expressed as:

$$\frac{\mathrm{d}[P_i]}{\mathrm{d}t} = \alpha_{p,i}V - \lambda[P_i] \tag{2}$$

Where $[P_i]$ is the the concentration of the RNA product, $\alpha_{p,i}$ the joint production rate from transcription and translation in the absence of regulation, and $\lambda$ is the stable molecule dilution rate for the protein.

#### 2.2 dCas9 Complex Kinetics

The differential equation system for all models incudes a reaction for the binding of Cas9 and gRNA into an active complex. While there are many other factors that can play into CRISPR kinetics, as these are not anticipated as being limited factors or being manipulated with regards to the specific system under investigation. We retain the binding, however, as degradation rate of gRNA can be large with respect to the binding rate when expression levels of Cas9 and gRNA are low, as will be the case during later stages of safety switch operation. When sgRNA binds to Cas9 protein, the resulting complex is significantly more stable, and is not expected to experience a significant rate of dissociation. Complex formation may thus be described as the reaction:

$$P_{\text{dCas9}} + G_i \xrightarrow{k_{C_g}} C_{\text{dCas9},i} \tag{3}$$

Where $P_{\text{dCas9}}$ is the dCas9 protein, $G_i$ is a particular gRNA, $C_{\text{dCas9},i}$ is the complex formed by the binding of these two molecules and $k_{C_g}$ is the reaction rate constant. This complex is affected by the same stable molecule diffusion as proteins, thus the following dynamics describe dCas9 complex binding.

$$\frac{\mathrm{d}[P_{\text{dCas9}}]}{\mathrm{d}t} = \alpha_{p,\text{dCas9}}V - \lambda[P_{\text{dCas9}}] - \sum_i k_{C_g}[P_{\text{dCas9}}][G_i] \tag{4}$$

$$\frac{\mathrm{d}[G_i]}{\mathrm{d}t} = \alpha_{r,i}V - \delta_g[G_i] - k_{C_g}[P_{\text{dCas9}}][G_i] \tag{5}$$

$$\frac{\mathrm{d}[C_{\text{dCas9},i}]}{\mathrm{d}t} = k_{C_g}[P_{\text{dCas9}}][G_i] - \lambda[C_{\text{dCas9},i}] \tag{6}$$

#### 2.3 Control Mechanisms: Transcription Factor Regulation

The activity of gRNA-guided dCas9 transcription factors is analogous to other types of transcription factor regulation, and so we modeled the effect of regulation on gene production using the standard Hill-Langmuir equation (CITE). Specifically, given the models of production given above, we model transcriptional regulation as a modulation of the effective transcription rate $\alpha_{r,i}$ or joint transcription/translation rate, $\alpha_{p,i}$.

As the production of a given species, $i$ is repressed by the dCas9 complex, $C_{\text{dCas9},i}$, then we model the modulation of the production rate in terms of the concentration of the transcription factor as:

$$\alpha_{p,i} = \alpha_{p,i}^0 \frac{K_R}{(K_R)^n + [C_{\text{dCas9},i}]^n} \tag{7}$$

Where $\alpha_{p,i}^0$ is the maximal production rate, $n$ is the Hill coefficient, and $K_R$ is the Hill Repression constant.

When there are multiple binding sites their effect on the target is multiplicative, so that each unique binding site has its own Hill-Langmuir term.

$$\alpha_{p,i} = \alpha_{p,i}^0 \prod_i \frac{K_R}{(K_R)^n + [C_{\text{dCas9},i}]^n} \tag{8}$$

In the case of identical binding sites, where the same dCas9-gRNA complex concentration is used for all of the Hill-Langmuir terms, this can be simplified to:

$$\alpha_{p,i} = \alpha_{p,i}^0 \left( \frac{K_R}{(K_R)^n + [C_{\text{dCas9},i}]^n} \right)^m \tag{9}$$

where $m$ is the total number of identical binding sites.

In both of these cases, we assume no interference between the gRNA target sites (e.g. like that which could be caused do to overlapping binding sites). To incorporate this interference, the concentration of the dCas9-gRNA complex could be modulated to reflect the effective concentration.

### 3 Circuit Models

From these components, models for all possible repressible promoters can be generating by connecting the equations above together and repeating the Hill-Langmuir terms the desired number of times. Models for three such generated promoters are presented below, the single gRNA repressor, and the heterogeneous and identical four-site repressors. The full models for all other systems are in the supplementary materials.

##### 3.1 0 Target Sites

$$\frac{\mathrm{d}V_2}{\mathrm{d}t} = -\lambda V_2 \tag{10}$$

$$\frac{\mathrm{d}[P_{\text{dCas9}}]}{\mathrm{d}t} = \alpha_{p,P_{\text{dCas9}}} V_2 - \lambda [P_{\text{dCas9}}] \tag{11}$$

$$\frac{\mathrm{d}[P_{\text{GFP}}]}{\mathrm{d}t} = \alpha_{p,P_{\text{GFP}}} V_2 - \lambda [P_{\text{GFP}}] \tag{12}$$

##### 3.2 1 Target Site

$$\frac{\mathrm{d}V_2}{\mathrm{d}t} = -\lambda V_2 \tag{13}$$

$$\frac{\mathrm{d}[G_1]}{\mathrm{d}t} = \alpha_{r,G_1} V_1 - k_{C_g}[G_1][P_{\text{dCas9}}] - \delta_g [G_1] \tag{14}$$

$$\frac{\mathrm{d}[C_{\text{dCas9},1}]}{\mathrm{d}t} = k_{C_g}[G_1][P_{\text{dCas9}}] - \lambda [C_{\text{dCas9},1}] \tag{15}$$

$$\frac{\mathrm{d}[P_{\text{dCas9}}]}{\mathrm{d}t} = \alpha_{p,P_{\text{dCas9}}} V_2 - \lambda [P_{\text{dCas9}}] - k_{C_g}[G_1][P_{\text{dCas9}}] \tag{16}$$

$$\frac{\mathrm{d}[P_{\text{GFP}}]}{\mathrm{d}t} = \alpha_{p,P_{\text{GFP}}} \frac{(K_R)^n}{(K_R)^n + [C_{\text{dCas9},1}]^n} V_2 - \lambda [P_{\text{GFP}}] \tag{17}$$

$$\frac{\mathrm{d}V_1}{\mathrm{d}t} = -\lambda V_1 \tag{18}$$

##### 3.3 2 Heterogeneous Target Sites

$$\frac{\mathrm{d}V_2}{\mathrm{d}t} = -\lambda V_2 \tag{19}$$

$$\frac{\mathrm{d}[G_1]}{\mathrm{d}t} = \alpha_{r,G_1} V_1 - k_{C_g}[G_1][P_{\text{dCas9}}] - \delta_g [G_1] \tag{20}$$

$$\frac{\mathrm{d}[C_{\text{dCas9},1}]}{\mathrm{d}t} = k_{C_g}[G_1][P_{\text{dCas9}}] - \lambda [C_{\text{dCas9},1}] \tag{21}$$

$$\frac{\mathrm{d}[G_2]}{\mathrm{d}t} = \alpha_{r,G_2} V_1 - k_{C_g}[G_2][P_{\text{dCas9}}] - \delta_g [G_2] \tag{22}$$

$$\frac{\mathrm{d}[C_{\text{dCas9},2}]}{\mathrm{d}t} = k_{C_g}[G_2][P_{\text{dCas9}}] - \lambda [C_{\text{dCas9},2}] \tag{23}$$

$$\frac{\mathrm{d}[P_{\text{dCas9}}]}{\mathrm{d}t} = \alpha_{p,P_{\text{dCas9}}} V_2 - \lambda [P_{\text{dCas9}}] - k_{C_g}[G_1][P_{\text{dCas9}}] - k_{C_g}[G_2][P_{\text{dCas9}}] \tag{24}$$

$$\frac{\mathrm{d}[P_{\text{GFP}}]}{\mathrm{d}t} = \alpha_{p,P_{\text{GFP}}} \frac{(K_R)^n}{(K_R)^n + [C_{\text{dCas9},1}]^n} \frac{(K_R)^n}{(K_R)^n + [C_{\text{dCas9},2}]^n} V_2 - \lambda [P_{\text{GFP}}] \tag{25}$$

$$\frac{\mathrm{d}V_1}{\mathrm{d}t} = -\lambda V_1 \tag{26}$$

##### 3.4 2 Identical Target Sites

$$\frac{\mathrm{d}V_2}{\mathrm{d}t} = -\lambda V_2 \tag{27}$$

$$\frac{\mathrm{d}[G_1]}{\mathrm{d}t} = \alpha_{r,G_1} V_1 - k_{C_g}[G_1][P_{\text{dCas9}}] - \delta_g[G_1] \tag{28}$$

$$\frac{\mathrm{d}[C_{\text{dCas9},1}]}{\mathrm{d}t} = k_{C_g}[G_1][P_{\text{dCas9}}] - \lambda[C_{\text{dCas9},1}] \tag{29}$$

$$\frac{\mathrm{d}[P_{\text{dCas9}}]}{\mathrm{d}t} = \alpha_{p,P_{\text{dCas9}}} V_2 - \lambda[P_{\text{dCas9}}] - k_{C_g}[G_1][P_{\text{dCas9}}] \tag{30}$$

$$\frac{\mathrm{d}[P_{\text{GFP}}]}{\mathrm{d}t} = \alpha_{p,P_{\text{GFP}}} \frac{(K_R)^n}{(K_R)^n + [C_{\text{dCas9},1}]^n} \frac{(K_R)^n}{(K_R)^n + [C_{\text{dCas9},1}]^n} V_2 - \lambda[P_{\text{GFP}}] \tag{31}$$

$$\frac{\mathrm{d}V_1}{\mathrm{d}t} = -\lambda V_1 \tag{32}$$

##### 3.5 3 Heterogeneous Target Sites

$$\frac{\mathrm{d}V_2}{\mathrm{d}t} = -\lambda V_2 \tag{33}$$

$$\frac{\mathrm{d}[G_1]}{\mathrm{d}t} = \alpha_{r,G_1} V_1 - k_{C_g}[G_1][P_{\text{dCas9}}] - \delta_g[G_1] \tag{34}$$

$$\frac{\mathrm{d}[C_{\text{dCas9},1}]}{\mathrm{d}t} = k_{C_g}[G_1][P_{\text{dCas9}}] - \lambda[C_{\text{dCas9},1}] \tag{35}$$

$$\frac{\mathrm{d}[G_2]}{\mathrm{d}t} = \alpha_{r,G_2} V_1 - k_{C_g}[G_2][P_{\text{dCas9}}] - \delta_g[G_2] \tag{36}$$

$$\frac{\mathrm{d}[C_{\text{dCas9},2}]}{\mathrm{d}t} = k_{C_g}[G_2][P_{\text{dCas9}}] - \lambda[C_{\text{dCas9},2}] \tag{37}$$

$$\frac{\mathrm{d}[G_3]}{\mathrm{d}t} = \alpha_{r,G_3} V_1 - k_{C_g}[G_3][P_{\text{dCas9}}] - \delta_g[G_3] \tag{38}$$

$$\frac{\mathrm{d}[C_{\text{dCas9},3}]}{\mathrm{d}t} = k_{C_g}[G_3][P_{\text{dCas9}}] - \lambda[C_{\text{dCas9},3}] \tag{39}$$

$$\frac{\mathrm{d}[P_{\text{dCas9}}]}{\mathrm{d}t} = \alpha_{p,P_{\text{dCas9}}} V_2 - \lambda[P_{\text{dCas9}}] - k_{C_g}[G_1][P_{\text{dCas9}}] - k_{C_g}[G_2][P_{\text{dCas9}}] - k_{C_g}[G_3][P_{\text{dCas9}}] \tag{40}$$

$$\frac{\mathrm{d}[P_{\text{GFP}}]}{\mathrm{d}t} = \alpha_{p,P_{\text{GFP}}} \frac{(K_R)^n}{(K_R)^n + [C_{\text{dCas9},1}]^n} \frac{(K_R)^n}{(K_R)^n + [C_{\text{dCas9},2}]^n} \frac{(K_R)^n}{(K_R)^n + [C_{\text{dCas9},3}]^n} V_2 - \lambda[P_{\text{GFP}}] \tag{41}$$

$$\frac{\mathrm{d}V_1}{\mathrm{d}t} = -\lambda V_1 \tag{42}$$

##### 3.6 3 Identical Target Sites

$$\frac{\mathrm{d}V_2}{\mathrm{d}t} = -\lambda V_2 \tag{43}$$

$$\frac{\mathrm{d}[G_1]}{\mathrm{d}t} = \alpha_{r,G_1} V_1 - k_{C_g}[G_1][P_{\text{dCas9}}] - \delta_g[G_1] \tag{44}$$

$$\frac{\mathrm{d}[C_{\text{dCas9},1}]}{\mathrm{d}t} = k_{C_g}[G_1][P_{\text{dCas9}}] - \lambda[C_{\text{dCas9},1}] \tag{45}$$

$$\frac{\mathrm{d}[P_{\text{dCas9}}]}{\mathrm{d}t} = \alpha_{p,P_{\text{dCas9}}} V_2 - \lambda[P_{\text{dCas9}}] - k_{C_g}[G_1][P_{\text{dCas9}}] \tag{46}$$

$$\frac{\mathrm{d}[P_{\text{GFP}}]}{\mathrm{d}t} = \alpha_{p,P_{\text{GFP}}} \frac{(K_R)^n}{(K_R)^n + [C_{\text{dCas9},1}]^n} \frac{(K_R)^n}{(K_R)^n + [C_{\text{dCas9},1}]^n} \frac{(K_R)^n}{(K_R)^n + [C_{\text{dCas9},1}]^n} V_2 - \lambda[P_{\text{GFP}}] \tag{47}$$

$$\frac{\mathrm{d}V_1}{\mathrm{d}t} = -\lambda V_1 \tag{48}$$

#### 3.7 4 Heterogeneous Target Sites

$$\frac{\mathrm{d}V_2}{\mathrm{d}t} = -\lambda V_2 \tag{49}$$

$$\frac{\mathrm{d}[G_1]}{\mathrm{d}t} = \alpha_{r,G_1} V_1 - k_{C_g}[G_1][P_{\text{dCas9}}] - \delta_g[G_1] \tag{50}$$

$$\frac{\mathrm{d}[C_{\text{dCas9},1}]}{\mathrm{d}t} = k_{C_g}[G_1][P_{\text{dCas9}}] - \lambda[C_{\text{dCas9},1}] \tag{51}$$

$$\frac{\mathrm{d}[G_2]}{\mathrm{d}t} = \alpha_{r,G_2} V_1 - k_{C_g}[G_2][P_{\text{dCas9}}] - \delta_g[G_2] \tag{52}$$

$$\frac{\mathrm{d}[C_{\text{dCas9},2}]}{\mathrm{d}t} = k_{C_g}[G_2][P_{\text{dCas9}}] - \lambda[C_{\text{dCas9},2}] \tag{53}$$

$$\frac{\mathrm{d}[G_3]}{\mathrm{d}t} = \alpha_{r,G_3} V_1 - k_{C_g}[G_3][P_{\text{dCas9}}] - \delta_g[G_3] \tag{54}$$

$$\frac{\mathrm{d}[C_{\text{dCas9},3}]}{\mathrm{d}t} = k_{C_g}[G_3][P_{\text{dCas9}}] - \lambda[C_{\text{dCas9},3}] \tag{55}$$

$$\frac{\mathrm{d}[G_4]}{\mathrm{d}t} = \alpha_{r,G_4} V_1 - k_{C_g}[G_4][P_{\text{dCas9}}] - \delta_g[G_4] \tag{56}$$

$$\frac{\mathrm{d}[C_{\text{dCas9},4}]}{\mathrm{d}t} = k_{C_g}[G_4][P_{\text{dCas9}}] - \lambda[C_{\text{dCas9},4}] \tag{57}$$

$$\frac{\mathrm{d}[P_{\text{dCas9}}]}{\mathrm{d}t} = \alpha_{p,P_{\text{dCas9}}} V_2 - \lambda[P_{\text{dCas9}}] - k_{C_g}[G_1][P_{\text{dCas9}}] - k_{C_g}[G_2][P_{\text{dCas9}}] - k_{C_g}[G_3][P_{\text{dCas9}}] - k_{C_g}[G_4][P_{\text{dCas9}}] \tag{58}$$

$$\frac{\mathrm{d}[P_{\text{GFP}}]}{\mathrm{d}t} = \alpha_{p,P_{\text{GFP}}} \frac{(K_R)^n}{(K_R)^n + [C_{\text{dCas9},1}]^n} \frac{(K_R)^n}{(K_R)^n + [C_{\text{dCas9},2}]^n} \frac{(K_R)^n}{(K_R)^n + [C_{\text{dCas9},3}]^n} \frac{(K_R)^n}{(K_R)^n + [C_{\text{dCas9},4}]^n} V_2 - \lambda[P_{\text{GFP}}] \tag{59}$$

$$\frac{\mathrm{d}V_1}{\mathrm{d}t} = -\lambda V_1 \tag{60}$$

#### 3.8 4 Identical Target Sites

$$\frac{\mathrm{d}V_2}{\mathrm{d}t} = -\lambda V_2 \tag{61}$$

$$\frac{\mathrm{d}[G_1]}{\mathrm{d}t} = \alpha_{r,G_1} V_1 - k_{C_g}[G_1][P_{\text{dCas9}}] - \delta_g[G_1] \tag{62}$$

$$\frac{\mathrm{d}[C_{\text{dCas9},1}]}{\mathrm{d}t} = k_{C_g}[G_1][P_{\text{dCas9}}] - \lambda[C_{\text{dCas9},1}] \tag{63}$$

$$\frac{\mathrm{d}[P_{\mathrm{dCas9}}]}{\mathrm{d}t} = \alpha_{p,P_{\mathrm{dCas9}}} V_2 - \lambda[P_{\mathrm{dCas9}}] - k_{C_g}[G_1][P_{\mathrm{dCas9}}] \tag{64}$$

$$\frac{\mathrm{d}[P_{\mathrm{GFP}}]}{\mathrm{d}t} = \alpha_{p,P_{\mathrm{GFP}}} \frac{(K_R)^n}{(K_R)^n + [C_{\mathrm{dCas9},1}]^n} \frac{(K_R)^n}{(K_R)^n + [C_{\mathrm{dCas9},1}]^n} \frac{(K_R)^n}{(K_R)^n + [C_{\mathrm{dCas9},1}]^n} \frac{(K_R)^n}{(K_R)^n + [C_{\mathrm{dCas9},1}]^n} V_2 - \lambda[P_{\mathrm{GFP}}] \tag{65}$$

$$\frac{\mathrm{d}V_1}{\mathrm{d}t} = -\lambda V_1 \tag{66}$$

#### 3.9 5 Heterogeneous Target Sites

$$\frac{\mathrm{d}V_2}{\mathrm{d}t} = -\lambda V_2 \tag{67}$$

$$\frac{\mathrm{d}[G_1]}{\mathrm{d}t} = \alpha_{r,G_1} V_1 - k_{C_g}[G_1][P_{\mathrm{dCas9}}] - \delta_g[G_1] \tag{68}$$

$$\frac{\mathrm{d}[C_{\mathrm{dCas9},1}]}{\mathrm{d}t} = k_{C_g}[G_1][P_{\mathrm{dCas9}}] - \lambda[C_{\mathrm{dCas9},1}] \tag{69}$$

$$\frac{\mathrm{d}[G_2]}{\mathrm{d}t} = \alpha_{r,G_2} V_1 - k_{C_g}[G_2][P_{\mathrm{dCas9}}] - \delta_g[G_2] \tag{70}$$

$$\frac{\mathrm{d}[C_{\mathrm{dCas9},2}]}{\mathrm{d}t} = k_{C_g}[G_2][P_{\mathrm{dCas9}}] - \lambda[C_{\mathrm{dCas9},2}] \tag{71}$$

$$\frac{\mathrm{d}[G_3]}{\mathrm{d}t} = \alpha_{r,G_3} V_1 - k_{C_g}[G_3][P_{\mathrm{dCas9}}] - \delta_g[G_3] \tag{72}$$

$$\frac{\mathrm{d}[C_{\mathrm{dCas9},3}]}{\mathrm{d}t} = k_{C_g}[G_3][P_{\mathrm{dCas9}}] - \lambda[C_{\mathrm{dCas9},3}] \tag{73}$$

$$\frac{\mathrm{d}[G_4]}{\mathrm{d}t} = \alpha_{r,G_4} V_1 - k_{C_g}[G_4][P_{\mathrm{dCas9}}] - \delta_g[G_4] \tag{74}$$

$$\frac{\mathrm{d}[C_{\mathrm{dCas9},4}]}{\mathrm{d}t} = k_{C_g}[G_4][P_{\mathrm{dCas9}}] - \lambda[C_{\mathrm{dCas9},4}] \tag{75}$$

$$\frac{\mathrm{d}[G_5]}{\mathrm{d}t} = \alpha_{r,G_5} V_1 - k_{C_g}[G_5][P_{\mathrm{dCas9}}] - \delta_g[G_5] \tag{76}$$

$$\frac{\mathrm{d}[C_{\mathrm{dCas9},5}]}{\mathrm{d}t} = k_{C_g}[G_5][P_{\mathrm{dCas9}}] - \lambda[C_{\mathrm{dCas9},5}] \tag{77}$$

$$\frac{\mathrm{d}[P_{\mathrm{dCas9}}]}{\mathrm{d}t} = \alpha_{p,P_{\mathrm{dCas9}}} V_2 - \lambda[P_{\mathrm{dCas9}}] - k_{C_g}[G_1][P_{\mathrm{dCas9}}] - k_{C_g}[G_2][P_{\mathrm{dCas9}}] - k_{C_g}[G_3][P_{\mathrm{dCas9}}] - k_{C_g}[G_4][P_{\mathrm{dCas9}}] - k_{C_g}[G_5][P_{\mathrm{dCas9}}] \tag{78}$$

$$\frac{\mathrm{d}[P_{\mathrm{GFP}}]}{\mathrm{d}t} = \alpha_{p,P_{\mathrm{GFP}}} \frac{(K_R)^n}{(K_R)^n + [C_{\mathrm{dCas9},1}]^n} \frac{(K_R)^n}{(K_R)^n + [C_{\mathrm{dCas9},2}]^n} \frac{(K_R)^n}{(K_R)^n + [C_{\mathrm{dCas9},3}]^n} \frac{(K_R)^n}{(K_R)^n + [C_{\mathrm{dCas9},4}]^n} \frac{(K_R)^n}{(K_R)^n + [C_{\mathrm{dCas9},5}]^n} V_2 - \lambda[P_{\mathrm{GFP}}] \tag{79}$$

$$\frac{\mathrm{d}V_1}{\mathrm{d}t} = -\lambda V_1 \tag{80}$$

#### 3.10 5 Identical Target Sites

$$\frac{\mathrm{d}V_2}{\mathrm{d}t} = -\lambda V_2 \tag{81}$$

$$\frac{\mathrm{d}[G_1]}{\mathrm{d}t} = \alpha_{r,G_1} V_1 - k_{C_g}[G_1][P_{\mathrm{dCas9}}] - \delta_g[G_1] \tag{82}$$

$$\frac{\mathrm{d}[C_{\mathrm{dCas9},1}]}{\mathrm{d}t} = k_{C_g}[G_1][P_{\mathrm{dCas9}}] - \lambda[C_{\mathrm{dCas9},1}] \tag{83}$$

$$\frac{\mathrm{d}[P_{\mathrm{dCas9}}]}{\mathrm{d}t} = \alpha_{p,P_{\mathrm{dCas9}}} V_2 - \lambda[P_{\mathrm{dCas9}}] - k_{C_g}[G_1][P_{\mathrm{dCas9}}] \tag{84}$$

$$\frac{\mathrm{d}[P_{\mathrm{GFP}}]}{\mathrm{d}t} = \alpha_{p,P_{\mathrm{GFP}}} \frac{(K_R)^n}{(K_R)^n + [C_{\mathrm{dCas9},1}]^n} \frac{(K_R)^n}{(K_R)^n + [C_{\mathrm{dCas9},1}]^n} \frac{(K_R)^n}{(K_R)^n + [C_{\mathrm{dCas9},1}]^n} \frac{(K_R)^n}{(K_R)^n + [C_{\mathrm{dCas9},1}]^n} \frac{(K_R)^n}{(K_R)^n + [C_{\mathrm{dCas9},1}]^n} V_2 - \lambda[P_{\mathrm{GFP}}] \tag{85}$$

$$\frac{\mathrm{d}V_1}{\mathrm{d}t} = -\lambda V_1 \tag{86}$$

#### 3.11 6 Heterogeneous Target Sites

$$\frac{\mathrm{d}V_2}{\mathrm{d}t} = -\lambda V_2 \tag{87}$$

$$\frac{\mathrm{d}[G_1]}{\mathrm{d}t} = \alpha_{r,G_1} V_1 - k_{C_g}[G_1][P_{\mathrm{dCas9}}] - \delta_g[G_1] \tag{88}$$

$$\frac{\mathrm{d}[C_{\mathrm{dCas9},1}]}{\mathrm{d}t} = k_{C_g}[G_1][P_{\mathrm{dCas9}}] - \lambda[C_{\mathrm{dCas9},1}] \tag{89}$$

$$\frac{\mathrm{d}[G_2]}{\mathrm{d}t} = \alpha_{r,G_2} V_1 - k_{C_g}[G_2][P_{\mathrm{dCas9}}] - \delta_g[G_2] \tag{90}$$

$$\frac{\mathrm{d}[C_{\mathrm{dCas9},2}]}{\mathrm{d}t} = k_{C_g}[G_2][P_{\mathrm{dCas9}}] - \lambda[C_{\mathrm{dCas9},2}] \tag{91}$$

$$\frac{\mathrm{d}[G_3]}{\mathrm{d}t} = \alpha_{r,G_3} V_1 - k_{C_g}[G_3][P_{\mathrm{dCas9}}] - \delta_g[G_3] \tag{92}$$

$$\frac{\mathrm{d}[C_{\mathrm{dCas9},3}]}{\mathrm{d}t} = k_{C_g}[G_3][P_{\mathrm{dCas9}}] - \lambda[C_{\mathrm{dCas9},3}] \tag{93}$$

$$\frac{\mathrm{d}[G_4]}{\mathrm{d}t} = \alpha_{r,G_4} V_1 - k_{C_g}[G_4][P_{\mathrm{dCas9}}] - \delta_g[G_4] \tag{94}$$

$$\frac{\mathrm{d}[C_{\mathrm{dCas9},4}]}{\mathrm{d}t} = k_{C_g}[G_4][P_{\mathrm{dCas9}}] - \lambda[C_{\mathrm{dCas9},4}] \tag{95}$$

$$\frac{\mathrm{d}[G_5]}{\mathrm{d}t} = \alpha_{r,G_5} V_1 - k_{C_g}[G_5][P_{\mathrm{dCas9}}] - \delta_g[G_5] \tag{96}$$

$$\frac{\mathrm{d}[C_{\mathrm{dCas9},5}]}{\mathrm{d}t} = k_{C_g}[G_5][P_{\mathrm{dCas9}}] - \lambda[C_{\mathrm{dCas9},5}] \tag{97}$$

$$\frac{\mathrm{d}[G_6]}{\mathrm{d}t} = \alpha_{r,G_6} V_1 - k_{C_g}[G_6][P_{\mathrm{dCas9}}] - \delta_g[G_6] \tag{98}$$

$$\frac{\mathrm{d}[C_{\mathrm{dCas9},6}]}{\mathrm{d}t} = k_{C_g}[G_6][P_{\mathrm{dCas9}}] - \lambda[C_{\mathrm{dCas9},6}] \tag{99}$$

$$\frac{\mathrm{d}[P_{\mathrm{dCas9}}]}{\mathrm{d}t} = \alpha_{p,P_{\mathrm{dCas9}}} V_2 - \lambda[P_{\mathrm{dCas9}}] - k_{C_g}[G_1][P_{\mathrm{dCas9}}] - k_{C_g}[G_2][P_{\mathrm{dCas9}}] - k_{C_g}[G_3][P_{\mathrm{dCas9}}] - k_{C_g}[G_4][P_{\mathrm{dCas9}}] - k_{C_g}[G_5][P_{\mathrm{dCas9}}] - k_{C_g}[G_6][P_{\mathrm{dCas9}}] \tag{100}$$

$$\frac{\mathrm{d}[P_{\mathrm{GFP}}]}{\mathrm{d}t} = \alpha_{p,P_{\mathrm{GFP}}} \frac{(K_R)^n}{(K_R)^n + [C_{\mathrm{dCas9},1}]^n} \frac{(K_R)^n}{(K_R)^n + [C_{\mathrm{dCas9},2}]^n} \frac{(K_R)^n}{(K_R)^n + [C_{\mathrm{dCas9},3}]^n} \frac{(K_R)^n}{(K_R)^n + [C_{\mathrm{dCas9},4}]^n} \frac{(K_R)^n}{(K_R)^n + [C_{\mathrm{dCas9},5}]^n} \frac{(K_R)^n}{(K_R)^n + [C_{\mathrm{dCas9},6}]^n} V_2 - \lambda[P_{\mathrm{GFP}}] \tag{101}$$

$$\frac{\mathrm{d}V_1}{\mathrm{d}t} = -\lambda V_1 \tag{102}$$

#### 3.12 6 Identical Target Sites

$$\frac{\mathrm{d}V_2}{\mathrm{d}t} = -\lambda V_2 \tag{103}$$

$$\frac{\mathrm{d}[G_1]}{\mathrm{d}t} = \alpha_{r,G_1} V_1 - k_{C_g}[G_1][P_{\mathrm{dCas9}}] - \delta_g [G_1] \tag{104}$$

$$\frac{\mathrm{d}[C_{\mathrm{dCas9},1}]}{\mathrm{d}t} = k_{C_g}[G_1][P_{\mathrm{dCas9}}] - \lambda [C_{\mathrm{dCas9},1}] \tag{105}$$

$$\frac{\mathrm{d}[P_{\mathrm{dCas9}}]}{\mathrm{d}t} = \alpha_{p,P_{\mathrm{dCas9}}} V_2 - \lambda [P_{\mathrm{dCas9}}] - k_{C_g}[G_1][P_{\mathrm{dCas9}}] \tag{106}$$

$$\frac{\mathrm{d}[P_{\mathrm{GFP}}]}{\mathrm{d}t} = \alpha_{p,P_{\mathrm{GFP}}} \frac{(K_R)^n}{(K_R)^n + [C_{\mathrm{dCas9},1}]^n} \frac{(K_R)^n}{(K_R)^n + [C_{\mathrm{dCas9},1}]^n} \frac{(K_R)^n}{(K_R)^n + [C_{\mathrm{dCas9},1}]^n} \frac{(K_R)^n}{(K_R)^n + [C_{\mathrm{dCas9},1}]^n} \frac{(K_R)^n}{(K_R)^n + [C_{\mathrm{dCas9},1}]^n} \frac{(K_R)^n}{(K_R)^n + [C_{\mathrm{dCas9},1}]^n} V_2 - \lambda [P_{\mathrm{GFP}}] \tag{107}$$

$$\frac{\mathrm{d}V_1}{\mathrm{d}t} = -\lambda V_1 \tag{108}$$

### 4 Parameter Fitting

Parameters needed for simulation were either taken from previously published work on CRISPR systems [2, 1], fit to data, or hypothesized based on observed repression in the lab. The complete set of parameter base values and their origin is listed in Table 2.

| Parameter | Meaning | Value | Source |
|---|---|---|---|
| $\alpha_{r,[1-6]}$ | Transcription rate, for $G_1 - G_6$ | $10^{3.3090}$ molecules/hour | [1] |
| $\alpha_{p,\text{GFP}}$ | Coupled transcription and translation rate, for $P_{\text{GFP}}$ | $10^{5.5793}$ molecules/hour | Fit to data |
| $\alpha_{p,\text{dCas9}}$ | Coupled transcription and translation rate, for $P_{\text{dCas9}}$ | $10^{2.0415}$ molecules/hour | [1] |
| $\delta_g$ | gRNA degradation rate | $10^{0.0003}$ fraction/hour | [1] |
| $\lambda$ | Stable Molecule (Protein and Plasmid) Dilution Rate | $10^{-1.6225}$ fraction/hour | Fit to data |
| $k_{C_g}$ | dCas9 and gRNA binding rate | $10^{-4.2577}$ | [1] |
| $n$ | Hill coefficient | 0.92 | [2] |
| $K_R$ | Hill Repression Constant | $10^{2.9}$ | Hypothesized |
| - | Initial Delay | 18.0149 Hours | Fit to data |

Table 2: Table of parameter symbols, meanings, values, and sources

The coupled transcription/translation rate of GFP, the stable molecule diffusion, and an initial lag, were all fit to an approximately 200 hour time course of Fluorolog-3 scanning fluorometer measurements that were normalized to Molecules of Equivalent Fluorescine (MEFL) units using flow cytometry measurements taken at the final time point. Parameters were fit using a least-squares fit for parameters in a logarithmic parameter calue space in MATLAB using the ODE for the ”No gRNA control” model.

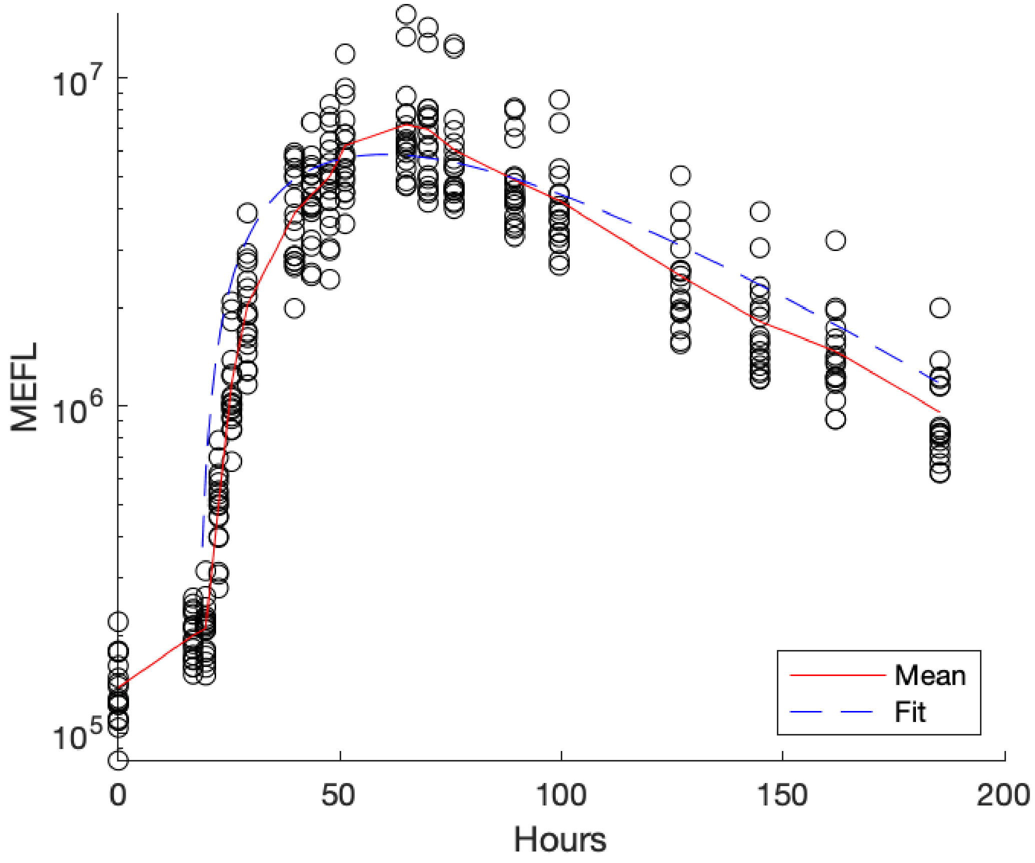

Figure 1: Experimental data and model simulation results after model fitting. A 200 hour time course of Fluorolog-3 scanning fluorometer measurements of GFP were converted to Molecules of Equivalent Fluorescine (MEFL) using a single flow cytometry measurement at the final time point. The individual data points (black circles) were used to the fit $\alpha_{p,\mathrm{GFP}}$, $\lambda$ and the Initial Delay. The results of a simulation of the "No gRNA control" model with these parameters is shown in blue. The mean value of the data points (red line) was not used directly, but is included to aid in comparison between the data and the simulation results.

### 5 Analysis of Repression Sensitivity

#### 5.1 Sensitivity to Parameter Values

To investigate the sensitivity of each promoter architecture to each individual parameter in the model, we modulated 6 parameters while holding all other values the same. Figure 2 shows the results of these perturbations for the 1 target site model, the 4 heterogeneous target site model, and the 4 identical target site model. Each of the parameters shown in figure 2 were modulated up and down by 2 standard deviations with a 1.5-fold log-normal uncertainty, except for the stable molecule diffusion rate ($\lambda$) which was modulated down 2 standard deviations, but was only modulated up 0.75 standard deviations, since any more would cause numerical instability.

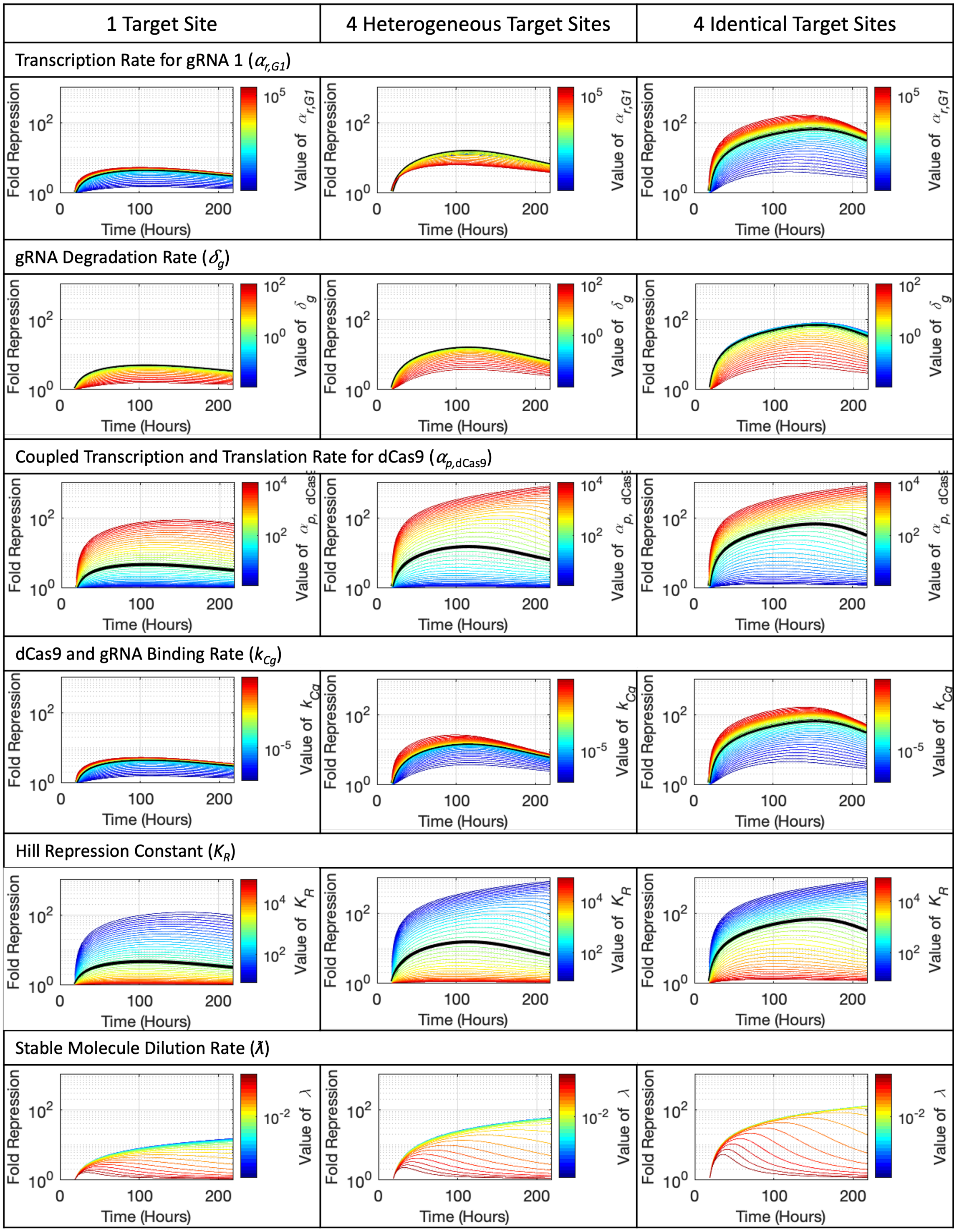

Figure 2: Examples of perturbation response for modulating individual parameters in 1 target site model, the 4 heterogeneous target site model, and the 4 identical target site model.

The modulation of most parameters show a monotonic trend, so that the lowest values (blue line) is on the opposite side of the trend of the highest value (red line). This was not the case for the $\alpha_{r,G_1}$ in the heterogenous target site models, including the 4 Heterogenous Target Site model (Figure 2 top center). In the case of this model, modulating the transcription rate of a single gRNA causes there to a different relative abundance of gRNAs in the total gRNA pool, but the total quantity of gRNA dow not become limiting because other dCas9-gRNA complexes are able to cause repression. For this reason, despite the larger range of possible fold repression values for the 4 identical target site model (Figure 2 top right) that model is not actually more sensitive to changes in the parameter value.

The remaining parameters showed similar trends for all three models, often with a similar range of fold repression values for the 4 identical target site model and 4 heterogeneous target site model, indicating that the identical target site models are not more sensitive to parameter values.

##### 5.1.1 Multi-Parameter Perturbations

Beyond the single parameter perturbations studies, we also performed an analysis in which all parameters were simultaneously and independently perturbed. For these multi-parameter perturbations, each parameter's base value was multiplied by a log-normal random factor drawn with a standard deviation of 1.1-fold (with $\pm$ 2 standard deviations spanning a range of just under 1.5-fold). Figure 3 shows the distribution of the trajectories from 10,000 simulations of each promoter architecture, where in each simulation the model parameters have been randomly and independently perturbed. A yellow color indicates a high number of simulations have passed through that given bin, where a dark blue color indicates that no simulations have. As with the single parameter perturbations, the range of GFP values remains comparable across the different promoters, and we see no marked difference between the results from perturbation between the heterogeneous and identical target site promoters.

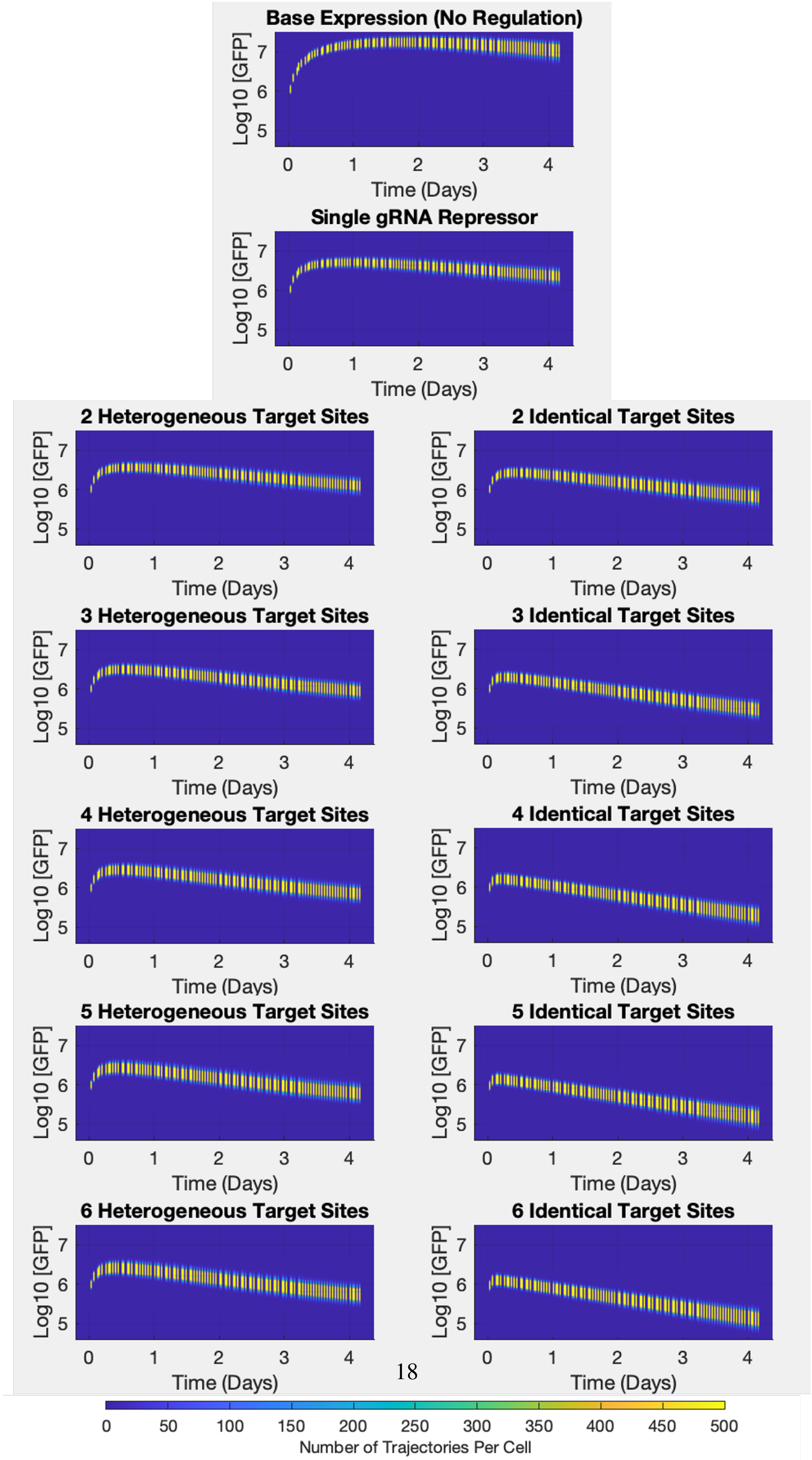

Figure 3: Distribution of trajectories generated by random perturbation of all parameters for all models.

### References

[1] Helen Scott, Dashan Sun, Jacob Beal, and Samira Kiani. Simulation-Based engineering of Time-Delayed safety switches for safer gene therapies. *ACS Synth. Biol.*, 11(5):1782–1789, May 2022.

[2] Junmin Wang, Samuel A Isaacson, and Calin Belta. Modeling genetic circuit behavior in transiently transfected mammalian cells. *ACS Synth. Biol.*, 8(4):697–707, April 2019.
